# m^6^A RNA methylation impairs gene expression variability and reproductive thermotolerance in *Arabidopsis*

**DOI:** 10.1101/2022.03.25.485737

**Authors:** Ling Wang, Haiyan Zhuang, Wenwen Fan, Xia Zhang, Haihong Dong, Hongxing Yang, Jungnam Cho

**Affiliations:** Shanghai Key Laboratory of Plant Functional Genomics and Resources, Shanghai Chenshan Botanical Garden, Shanghai 201602, China; National Key Laboratory of Plant Molecular Genetics, CAS Center for Excellence in Molecular Plant Sciences, Shanghai Institute of Plant Physiology and Ecology, Chinese Academy of Sciences, Shanghai 200032, China; University of Chinese Academy of Science, Beijing 100049, China; CAS-JIC Centre for Excellence in Plant and Microbial Science, Shanghai 200032, China

**Keywords:** m^6^A RNA methylation, gene expression variability, heat tolerance, *AtALKBH10B*, *Arabidopsis thaliana*

## Abstract

Plants are more susceptible to high temperature stress during reproductive development, which can cause drastic yield loss of fruit and seed crops. Unfortunately, the underlying mechanism remains largely unknown. Here we suggest that m^6^A RNA methylation level increases in the reproductive tissues of *Arabidopsis* and negatively regulates gene expression variability. It has been suggested that stochasticity of gene expression can be advantageous to fitness of living organisms under environmental challenges. Indeed, reduced gene expression variability in flowers was associated with compromised transcriptional activation of heat-responsive genes. Importantly, disruption of an RNA demethylase *AtALKBH10B* led to lower gene expression variability, hypo-responsiveness of heat-activated genes, and strong reduction of plant fertility. Overall, our work proposes a novel mechanism that m^6^A RNA modification mediates the bet-hedging strategy of plants challenged by heat stress.

## Introduction

Plant reproductive organs are particularly more susceptible to environmental challenges such as heat stress (Chaturvedi et al., 2021; Jacott and Boden, 2020). It has been well documented that in the heat stress response during the reproductive development, genes involved in the unfolded protein response (UPR) play an important role (Deng et al., 2011; Gao et al., 2021; Iwata and Koizumi, 2005; Zhang et al., 2017). Two basic leucine-zipper domain containing transcription factors, bZIP28 and bZIP60, are essential for maintaining fertility under heat stress (Deng et al., 2011; Zhang et al., 2017). More recently, a paralog of bZIP28, bZIP17, was also suggested to play a role in the reproductive heat tolerance in *Arabidopsis* (Gao et al., 2021). In addition, SQUAMOSA PROMOTER BINDING PROTEIN-LIKE (SPL) transcription factors, SPL1 and SPL12, are also critical genetic elements required for thermotolerance at the reproductive stage of *Arabidopsis* (Chao et al., 2017). In addition to the DNA level of control mediated by transcription factors, plants’ reproductive thermos-sensitivity can be regulated at post-transcriptional step. For example, a number of small RNAs including small interfering (si) RNAs and micro (mi) RNAs were identified to be responsive to heat stress at the reproductive stages of flax, soybean and maize (Ding et al., 2021; He et al., 2019; Pokhrel and Meyers, 2022). Overall, hypersensitivity to heat stress during the reproductive development is under complex regulation involving various transcription factors and small RNAs.

Sporadic and random gene expression was considered a technical noise in transcriptome study; however, it has been becoming increasingly appreciated as an important cellular signal governing various biological process (Eldar and Elowitz, 2010; Grimbergen et al., 2015; de Jong et al., 2019). In unicellular organisms, stochastic and noisy transcriptional initiation is considered one of the strategies to increase fitness in adverse growth conditions (Elowitz et al., 2002; Grimbergen et al., 2015; Jones et al., 2014; Liu et al., 2015). Stochasticity of gene expression is also often observed in multicellular organisms; for example, clonal heterogeneity of gene expression is important for lineage choice of mouse haematopoietic progenitor cells (Chang et al., 2008), the intestinal differentiation of nematode is also influenced by gene expression variability (Raj et al., 2010), and a robust transcription of NF-κB is attributed to both intrinsic and extrinsic variability (Kellogg and Tay, 2015). In addition, several previous studies also investigated the variable gene expression behavior in *Arabidopsis* (Araújo et al., 2017; Bhosale et al., 2013; Cortijo et al., 2019; Hirao et al., 2015). These studies determined the gene expression variability from different scales ranging from cellular, organismal, and population levels. Of note, the hypervariable genes in expression were suggested to be enriched with factors involved in environmental responses (Cortijo et al., 2019; Hirao et al., 2015). Taken altogether, transcriptional noise is an evolutionary conserved strategy that enhances gene expression robustness, particularly in response to adverse conditions.

In this study, we investigated the relevance of gene expression variability in the reproductive thermos-sensitivity of *Arabidopsis*. We also link the noise in gene expression with a specific kind of RNA modification, N6-methyladenosine (m^6^A), which is the most widespread RNA modification present in mRNAs (Jia et al., 2013; Meyer and Jaffrey, 2017; Yue et al., 2019). Our work provides a novel insight into how RNA modification controls gene expression stochasticity and thereby heat tolerance, which overall can help mitigate the heat-triggered reproductive failure of agriculturally important crops.

## Results

### Developmentally divergent heat response in transcriptome and epitranscriptome

Plant response to environmental challenges varies during development (Chaturvedi et al., 2021; Jacott and Boden, 2020). For example, it is well known that reproductive tissues are more sensitive to heat stress (Jacott and Boden, 2020). Such heat susceptibility during the reproductive development can cause reduction of yield and quality of fruit and cereal crops. To dissect the molecular mechanisms underlying the heat-imposed reproductive failure in *Arabidopsis*, we carried out the transcriptomic analyses of the heat-stressed flower and leaf samples. As shown in Fig. 1a, flower exhibited a distinct pattern of gene expression under heat stress. Genes in cluster 1 and 10, for instance, were strongly activated upon heat treatment in leaves, while in flowers the level of heat activation was drastically reduced (Fig. 1a). On the other hand, genes in cluster 4 and 9 were hyper-activated under heat stress in flowers (Fig. 1a).

**Fig. 1.**
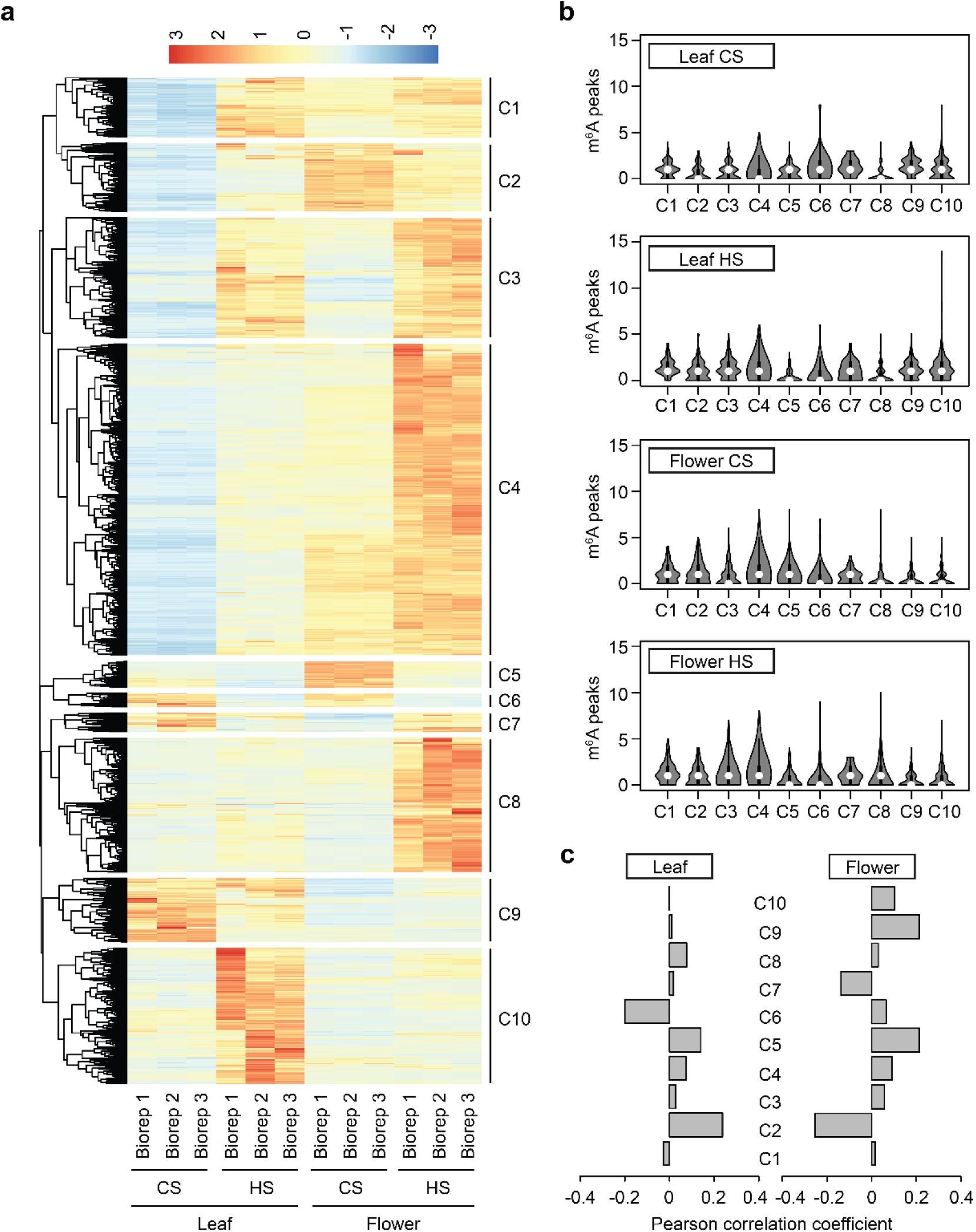
Divergent response to heat stress in leaf and flower. **a**. Heatmap of transcriptome data generated from the control and heat-stressed *Arabidopsis* leaves and flowers. Each row represents individual gene and genes are grouped to ten clusters by their expression pattern similarity. Genes with detectable level of expression are only considered (FPKM of at least 1 in any of the samples tested). CS, control sample; HS, heat-stressed sample; Biorep, biological replication. **b**. Violin plot displaying the distribution of the number of m^6^A peaks per transcript in different samples. **c**. Pearson correlation coefficient of fold change of expression and m^6^A enrichment in heat. The m^6^A enrichment was determined by the log2-fold change of m^6^A levels to input levels.

Previous studies suggested that RNA modifications (often referred to as epitranscriptomic marks) are implicated in cellular response to environmental challenges (Meyer et al., 2015; Scutenaire et al., 2018; Zhou et al., 2015); however, the functional roles of RNA modification in stress response remain largely obscure. In order to test if the developmentally divergent heat responsiveness can be attributed to m^6^A RNA methylation, the most widespread RNA modification type, we carried out the m^6^A-RNA immunoprecipitation (RIP)-seq experiments using the same samples tested in Fig. 1a. Figure 1b shows the number of m^6^A peaks per transcript in different samples presented in gene clusters defined in Fig. 1a. In leaves, cluster 2 and 4 showed slight increase in their m^6^A peak numbers under heat and the flower sample exhibited marginal increase of peak numbers in cluster 3 and 8 in the heat-stressed condition (Fig. 1b). In addition, we also measured the enrichment of m^6^A RNA methylation by comparing the levels of m^6^A-immunoprecipitated and total RNA. Of note, as shown in Fig. S1, the m^6^A enrichment showed very weak correlation with gene expression levels in all samples tested, suggesting that RNA methylation does not directly influence the mRNA levels. This is in strong agreement with previous studies that suggested a marginal effect of RNA methylation in mRNA steady-state levels (Anderson et al., 2018).

We then asked if m^6^A RNA modification might impact the heat responsiveness of gene expression. To test this, the fold change of gene expression under heat and difference of m^6^A enrichment between non-stressed and stressed samples was correlated. Figure 1c shows the Pearson correlation coefficient of gene expression and m^6^A enrichment changes under heat in both leaves and flowers. In both samples, we were not able to detect any meaningful levels of correlation between gene expression and m^6^A enrichment changes upon heat stress. Taken altogether, despite the vast changes of transcriptome profile under heat stress, m^6^A RNA methylation seems to have minimal changes under heat stress and thus limited impact on mRNA levels.

While investigating the m^6^A RNA methylation data, we found that the flower samples display strikingly stronger enrichment of m^6^A (Fig. 2a and b), which is also consistent with previous studies (Wan et al., 2015; Wang et al., 2022). As shown in Fig. 2a, the distribution of m^6^A mark in the flower samples was different from that of leaves exhibiting higher RNA methylation along the CDS. Figure 2b also shows that flowers contain more m^6^A-methylated transcripts. Similarly, the flower samples included more mRNAs that contain higher number of m^6^A peaks (Fig. 2c), which collectively suggest that m^6^A RNA methylation is developmentally controlled and undergoes minor changes upon heat stress.

**Fig. 2.**
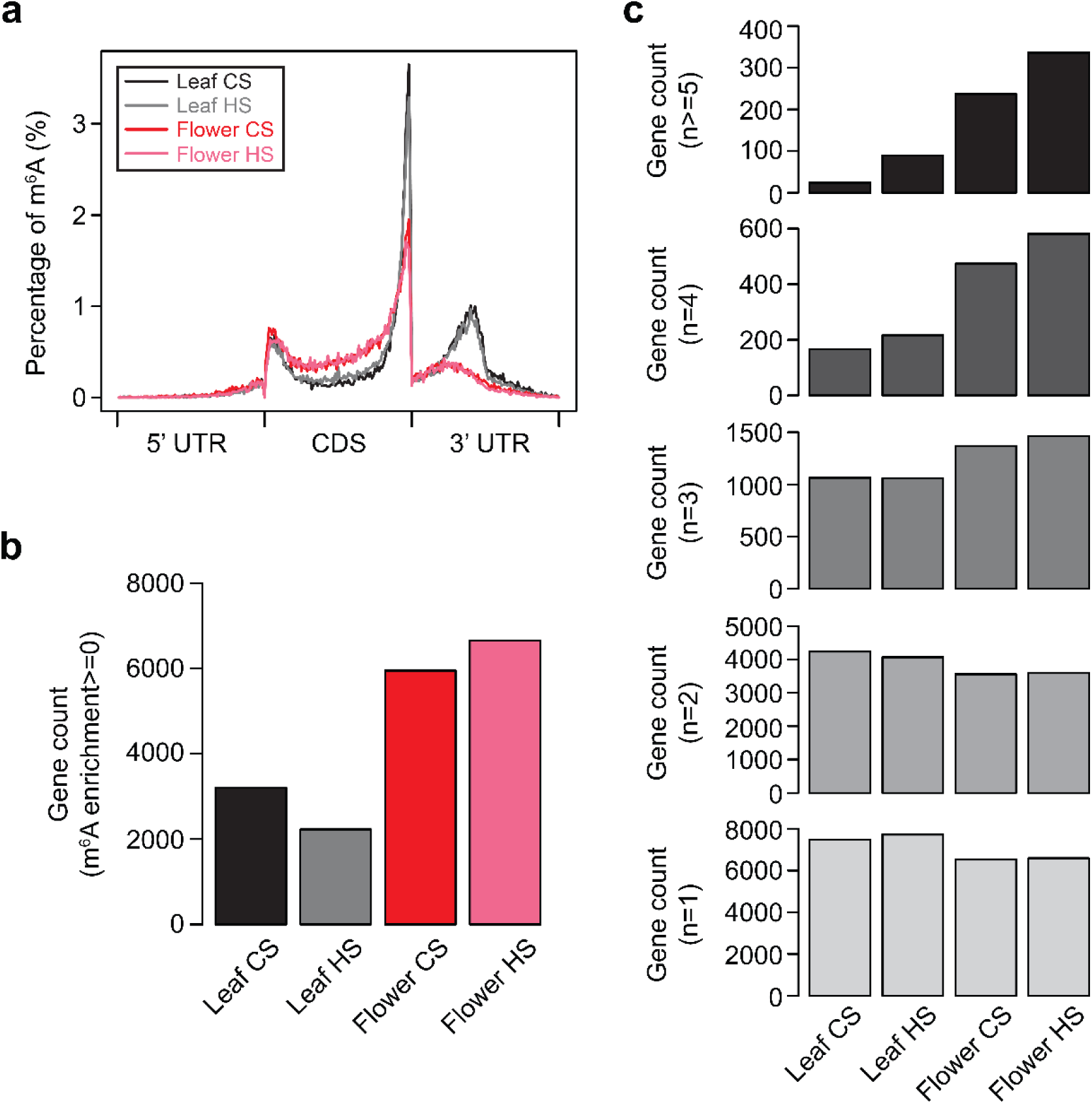
Increased RNA methylation in flowers. **a**. Distribution of m^6^A in 5’ UTR, CDS and 3’ UTR. Regions of individual transcript were divided to 100 non-overlapping windows to which the mid-point of each identified m^6^A peak was mapped and counted. **b**. Count of genes, the m^6^A enrichment score of which is above 0. The m^6^A enrichment was as defined in Fig. 1c. **c**. Count of genes that contain the m^6^A peaks for the indicated numbers (from bottom to top, 1 to 5).

### Reduced expression variability of m^6^A-marked transcripts

Our data so far suggested that flowers are distinct from leaves exhibiting stronger m^6^A RNA methylation which does not seem to directly contribute to divergent transcriptomic changes triggered by heat stress. We thus next investigated how m^6^A modification dictates gene expression dynamics in the reproductive thermal response. Intriguingly, we found a strong negative correlation between m^6^A enrichment and gene expression variability. Gene expression variability refers to stochastic and noisy expression of genetically identical cells and can be determined by the squared coefficient of variation (CV^2^). A previous report suggested that gene expression variability can be also observed and measured for a group of plants from which RNA was extracted from individual plants (Bhosale et al., 2013; Cortijo et al., 2019). In this study, we generated RNA-seq datasets for three independent biological replicates and interestingly, gene expression variability determined from biological replications was similar to what was calculated from single-cell (sc) transcriptome data (Fig. S2). This suggests that biological replications can be used to infer gene expression variability.

We then compared the gene expression variability of the transcripts containing varying number of m^6^A peaks and found a strong negative correlation (Fig. 3a and b). In both leaves and flowers, transcripts harboring a greater number of RNA methylation peaks showed drastically reduced gene expression variability (Fig. 3a and b). In addition, the number of lowly variable genes (LVGs) was higher in flowers (Fig. 3d and e), while highly variable genes (HVGs) were found similar in leaves and flowers (Fig. 3c). Interestingly, LVGs found in flowers were strongly enriched with genes involved in abiotic stress response (Fig. S3), which partly indicates a functional association between reduced expression variability and stress resistance. We also examined the HVGs and LVGs determined by other studies for their m^6^A RNA methylation levels. In the study of Cortijo et al., HVGs and LVGs were identified from inter-individual expression variability (Cortijo et al., 2019), and as shown in Fig. 3f, LVGs contained more m^6^A-methylated transcripts. Similar result was also observed for HVGs and LVGs identified from a scRNA-seq dataset. We analyzed a previously published scRNA-seq generated from *Arabidopsis* root tip (Zhang et al., 2019), and determined both HVGs and LVGs. Consistently, LVGs mostly contained at least one m^6^A peak, whereas HVGs exhibited very weak RNA methylation (Fig. 3g). In summary, the increased RNA methylation in flowers is associated with lower gene expression variability.

**Fig. 3.**
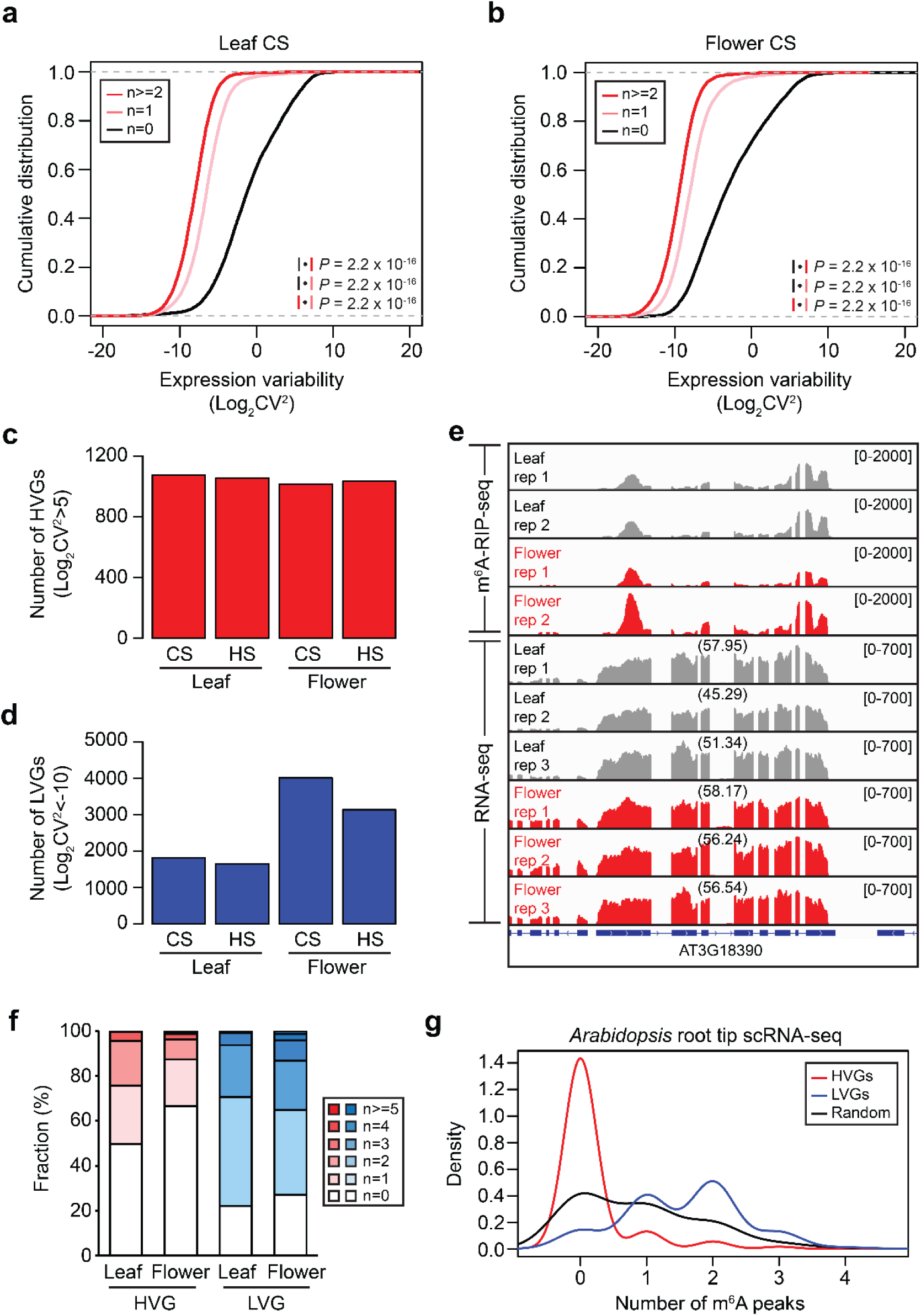
RNA methylation and expression variability is inversely correlated. **a** and **b**. Cumulative distribution of gene expression variability in leaf (**a**) and flower (**b**) for transcripts containing different number of m^6^A peaks. Gene expression variability was defined as log2-convered coefficient of variation (CV^2^). **c** and **d**. Number of HVGs (**c**) and LVGs (**d**) in different samples. HVG, highly variable gene; LVG, lowly variable gene. **e**. Genome browser snapshot for a representative locus exhibiting the increased m^6^A level and decreased expression variability in the flower tissue. RNA-seq data was generated from the same samples used for m^6^A-RIP-seq. Rep, biological replication. Numbers in parentheses indicate FPKM of corresponding gene. **f**. Fraction of genes with indicated number of m^6^A peaks. HVG and LVG was obtained from the study of Cortijo et al. **g**. Density distribution of m^6^A peak number per transcript. HVG and LVG was identified from single-cell RNA-seq data generated from *Arabidopsis* root tip.

### *AtALKBH10B* regulates expression variability and reproductive thermotolerance

In search for possible regulators of m^6^A-methylated transcriptome, we paid attention to *AtALKBH10B* (hereinafter *10B*) because its expression level is the highest among RNA demethylases and becomes higher in flower tissues (Fig. S4). To test if *10B* plays a role in the m^6^A-associated gene expression variability, we generated the m^6^A-RIP-seq data using the *10b-1* mutants. In the m^6^A distribution analyses, we were not able to see any significant differences between the wt and *10b-1* leaves (Fig. 4a). However, the flower samples of *10b-1* exhibited distinct m^6^A distribution as compared with wt, showing stronger peak around the stop codon (Fig. 4b). In addition, the m^6^A enrichment level was increased significantly in flowers of *10b-1*, while in leaves the difference of RNA methylation was only marginal (Fig. 4c). This altogether suggests that 10B mediates RNA demethylation mostly in flowers of *Arabidopsis*.

**Fig. 4.**
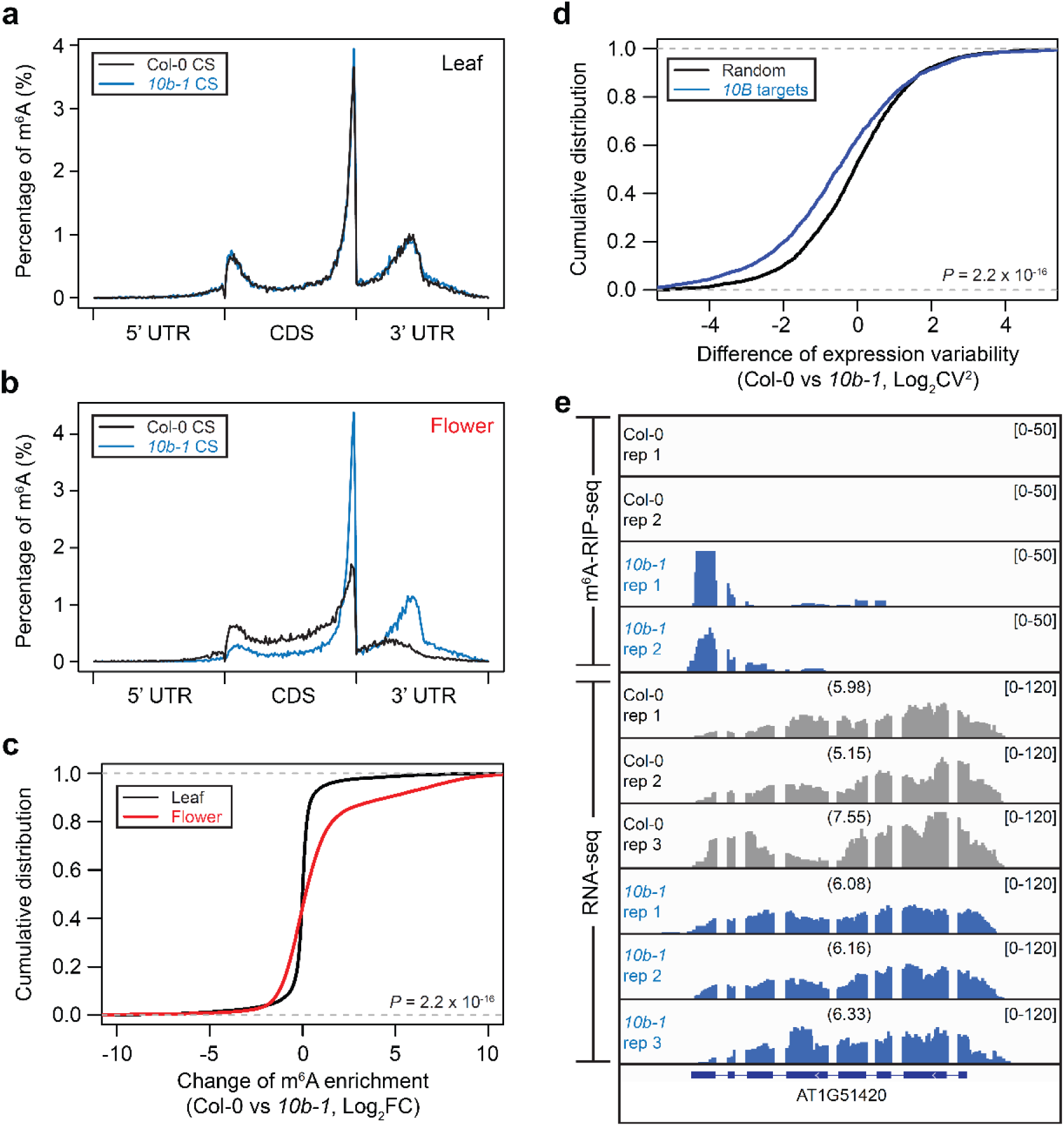
*AtALKBH10B* regulates m^6^A RNA methylation and expression variability. **a** and **b**. m^6^A distribution of wt and *10b-1* in leaf (**a**) and flower (**b**). **c**. Cumulative distribution of the m^6^A enrichment fold change in *10b-1* normalized against Col-0. *P* value was obtained by the one-tailed Wilcoxon rank sum test. **d**. Cumulative distribution of expression variability difference between wt and *10b-1* determined for the *10B*-regulated and randomly selected transcripts. The 10B target genes are those with at least two-fold of m^6^A enrichment compared to wt. *P* value was obtained by the one-tailed Wilcoxon rank sum test. **e**. Genome browser snapshot for a representative locus displaying a strong m^6^A enrichment and low expression variability in the *10b-1* mutant.

Next, we wanted to assess the gene expression variability of the *10B*-regulated genes. The transcripts with increased RNA methylation in the *10b-1* mutant was identified as *10B* targets. Gene enrichment analyses revealed that *10B* targets include genes encoding for RNA binding proteins and those involved in RNA regulation (Fig. S5), which is partly consistent to what was observed for LVGs of flowers (Fig. S3). The gene expression variability of the *10B* target genes were then compared with those of randomly selected genes. Interestingly, we found that the *10B*-targeted transcripts exhibit reduced level of gene expression variability (Fig. 5d and e). These data further support the notion that increased m^6^A RNA methylation leads to reduction in gene expression variability.

**Fig. 5.**
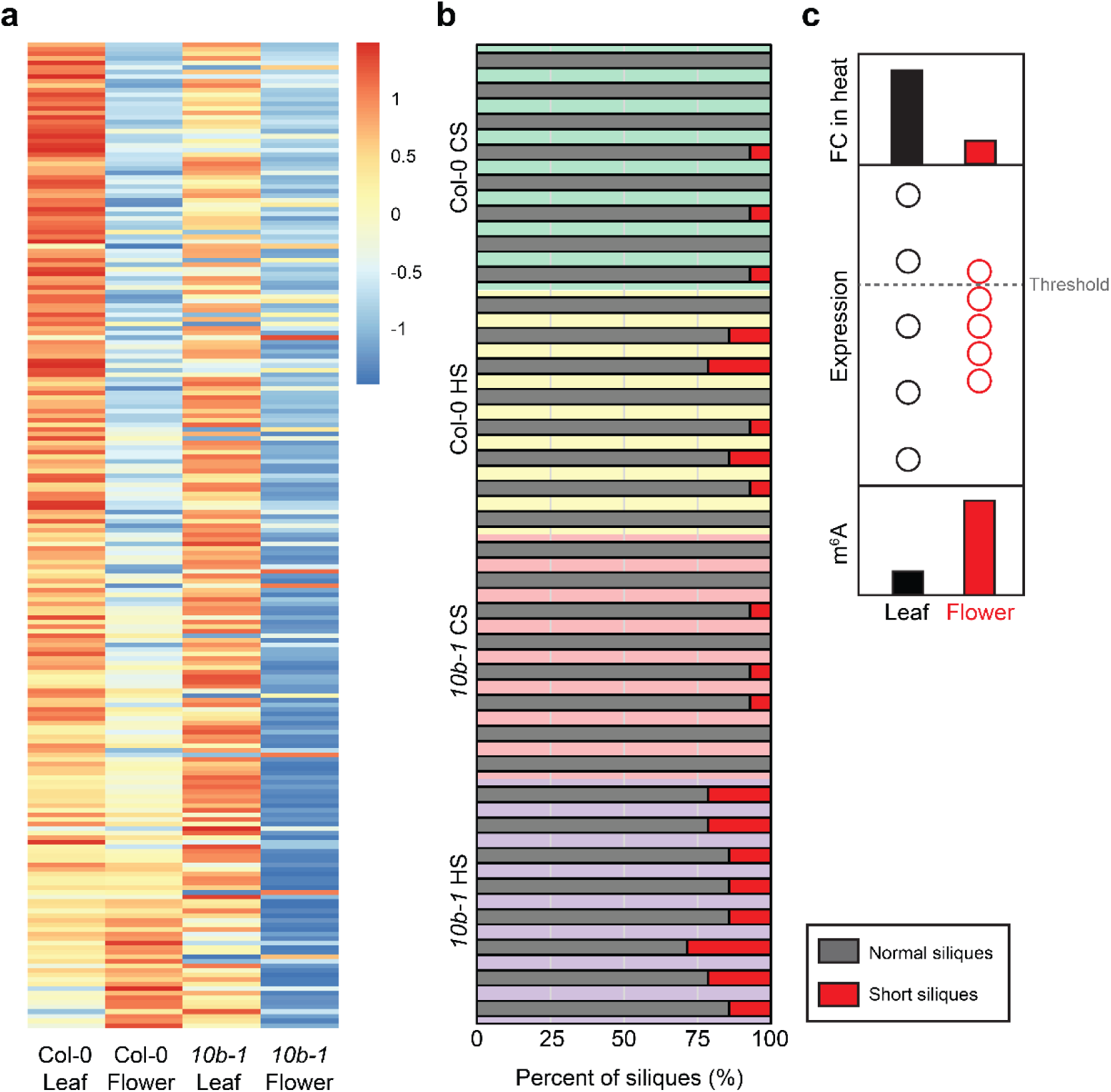
Reduced heat-responsiveness and heat tolerance of *10b* mutant flower. **a**. Heatmap of FC expression upon heat stress in different samples. **b**. Fraction of normal and short siliques of plants subjected to control and 6 h-heat stress condition. CS, control sample; HS, heat-stressed sample. Siliques that are longer than 10 mm were considered normal and those shorter than 10 mm were counted as short siliques. **c**. Schematic illustration of m^6^A-associated gene expression variability. Floral tissues exhibit stronger m^6^A RNA methylation, lower expression variability, and weaker heat-activation.

In order to understand the functional relevance of gene expression variability in the heat stress response of *Arabidopsis*, we examined the heat responsiveness of genes in leaves and flowers of wt and *10b-1*. For this, the heat-activated genes in the wt leaves were selected and their fold change expression in heat was compared. Noticeably, the fold change of heat-activated genes was drastically compromised in the wt flowers, and such reduction became more prominent in the flowers of *10b-1* mutant (Fig. 5a). Importantly, the heat-stressed *10b-1* mutant plants contained higher number of aberrant siliques, indicating that *10B* is required for heat resistance of *Arabidopsis* flowers. In conclusion, *10B* regulates m^6^A RNA methylation mainly in flowers and contributes to both variable gene expression and heat responsiveness, all of which is required for reproductive success under environmentally challenging condition.

## Discussion

It has been previously suggested that stochastic gene expression is advantageous to unicellular organisms with respect to survival under threatening environmental conditions (Cortijo et al., 2019; Elowitz et al., 2002; Grimbergen et al., 2015; Hirao et al., 2015). Such variable gene expression behavior was also observed in higher eukaryotes including mammals and was found to be associated with varying level of susceptibility to diverse diseases (Green et al., 2020; Hagai et al., 2018; de Jong et al., 2019). Gene expression variability can be determined at different levels ranging from cellular to population scales. Recently, Cortijo et al. investigated inter-individual gene expression variability of *Arabidopsis* during the diurnal circadian oscillation and discovered many HVGs involved in environmental responses (Cortijo et al., 2019). In this study, gene expression variability was determined from independent biological replications, and we showed that it was robust enough to represent cellular level of gene expression variability (Fig. S2). Importantly, gene expression became less variable in the reproductive tissues and this in turn caused reduced activation of heat-responsive genes and reproductive failure by heat stress (Fig. 5a and c). Therefore, our work demonstrates for the first time in a plant system that gene expression variability is a key to reproductive success in heat stress.

m^6^A RNA methylation has been relatively well characterized to be implicated in diverse RNA processing in plants (Chandola et al., 2015; Yue et al., 2019). It is worth mentioning that we showed in this study the genetic evidence that m^6^A RNA methylation negatively regulates gene expression variability (Fig. 3). Previously, several studies proposed that promoter structure, DNA methylation and chromatin status dictate stochastic gene transcription (Bashkeel et al., 2019; Cortijo et al., 2019; Jones et al., 2014). Further to the control at DNA or chromatin level, we suggest that an epitranscriptomic mark adds to the complex regulation of gene expression variability. Given that m^6^A-methylated mRNAs are preferably located to cytoplasmic stress granules (SGs) (Alvarado-Marchena et al., 2021; Fu and Zhuang, 2020; Ries et al., 2019; Scutenaire et al., 2018), we speculate that the physical confinement of m^6^A-methylated transcripts to SGs might play a role in stable gene expression variability. To test this possibility, we compared LVGs and random genes for their SG enrichment score determined in our previous study (Kim et al., 2021), and LVGs indeed showed stronger SG enrichment (Fig. S6). However, the precise mechanisms as to how low gene expression variability is attributed to SG localization require further investigation. Overall, our study unveils a hidden role for m^6^A RNA methylation in gene expression variability, accounting for heat stress sensitivity of reproductive organs of *Arabidopsis*.

## Materials and Methods

### Plant materials and growth condition

*Arabidopsis* Col-0 and *atalkbh10b-1* (SALK_004215C) mutant plants were grown at 22 °C under 16 h light/8 h dark day/night cycle. Heat stress treatment was performed to 5-week-old plants by elevating the growth temperature to 37 °C for 3 h. Rosette leaves (n=6) and stage 1 to 12 floral buds (n=12) were collected for both RNA-seq (3 biological replicates) and m^6^A-RIP-seq (2 biological replicates).

### NGS library construction

Total RNA was isolated using the Trizol reagent (Invitrogen). For m^6^A-RIP-seq, 50 μg of total RNA was used. Briefly, poly(A) RNA was selected using the oligo-d(T)25 magnetic beads (Thermo Fisher) and was fragmented using Magnesium RNA Fragmentation Module (NEB) at 86 °C for 7 min. Cleaved RNA fragments were then incubated for 2 h at 4 °C with m^6^A-specific antibody (No. 202003, Synaptic Systems) in the IP buffer (50 mM Tris-HCl, 750 mM NaCl and 0.5% Igepal CA-630). Library preparation was performed using the NEBNext Ultra Directional RNA Library Prep Kit following the manufacturer’s instructions. Finally, 150 bp paired-end sequencing (PE150) were carried out on an Illumina Novaseq 6000. RNA-seq libraries were constructed by following the same method.

### NGS data analyses

Raw sequencing data were cleaned using Trimmomatic (version 0.39) to remove reads containing adapter and low-quality sequences (Bolger et al., 2014). Trimmed reads were then aligned to the *Arabidopsis* reference genome (TAIR10) with default settings using Hisat2 (version 2.2.1) (Kim et al., 2015). FPKM values were calculated by StringTie (version 2.1.7) (Pertea et al., 2015). Visualization of the sequencing data was performed using the Integrative Genomics Viewer (IGV) (Robinson et al., 2011). For m^6^A peak calling, MACS2 (version 2.2.7.1) was run with the following parameters: --nomodel,--extsize 50, -p 5e-2, and -g 65084214 (Zhang et al., 2008). The -g option accounts for the size of the *Arabidopsis* transcriptome. m^6^A peaks identified in both two replicates were annotated by CHIPseeker (Yu et al., 2015). Distribution of m^6^A peaks along mRNAs were analyzed by adopting the MeRIP-PF scripts (Li et al., 2013).

### Heat resistance phenotyping

Heat tolerance test at reproductive stage was carried out as described previously (Zhang et al., 2017). Briefly, plants were grown under normal growth condition set at 22 °C and 16 h light/8 h dark day/night cycle. Heat stress of 37 °C was treated for 6 h when the first flower appeared. After the heat treatment, plants were moved to normal growth condition and continued to grow for 10 d. Silique length was measured from eight individual plants per genotype.

## Data availability

The NGS data generated in this study are deposited to SRA repository under PRJNA793364 and summarized in Supplementary Table 1. The analyses were performed using the standard codes instructed by the tools described in the Methods and the custom codes used in this study are deposited to GitHub (https://github.com/JungnamChoLab).

## Acknowledgements

This work was supported by the Strategic Priority Research Program of Chinese Academy of Sciences (XDB27030209), National Natural Science Foundation of China (31970518, 32150610473 and 32111540256) and General Program of Natural Science Foundation of Shanghai (22ZR1469100) granted to JC. HY is supported by the Shanghai Landscaping Administrative Bureau Program (G172404) and Shanghai Pujiang Program (16PJ1403000).

## Author contribution

LW, HZ and HY conceived the idea and designed the experiments. LW, HZ, WF, XZ and HD conducted the experiments. LW and JC analyzed the data. LW and JC wrote the manuscript. JC revised the manuscript.

