## Supplementary information for "m^6^A RNA methylation impairs gene expression variability and reproductive thermotolerance in *Arabidopsis*"

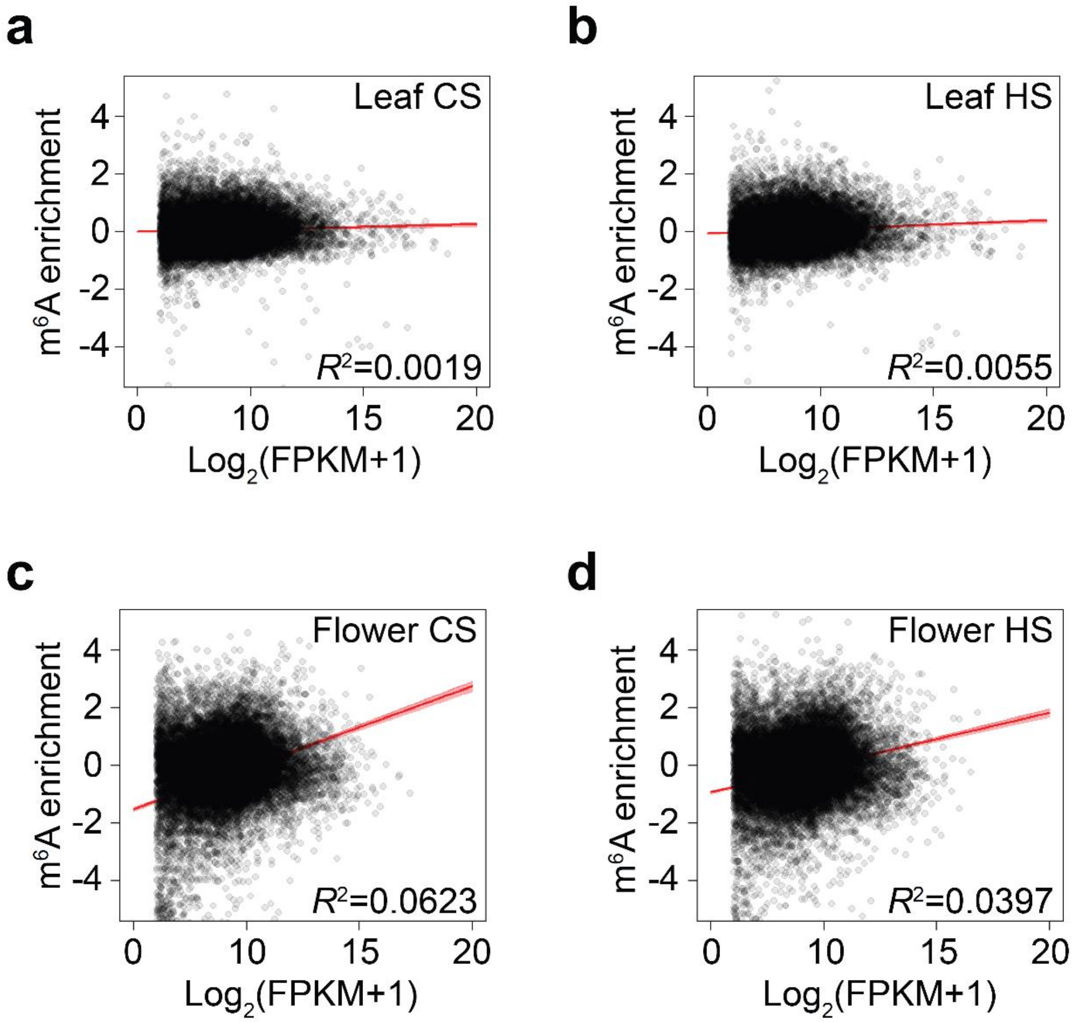


**Fig. S1 | Correlation between m$^6$A and expression levels.**

**a**-**d**. The enrichment of m$^6$A and log$_2$-converted FPKM values are plotted for leaves of non-stressed plants (**a**), heat-stressed leaves (**b**), non-stressed flowers (**c**), and heat-stressed flowers (**d**). Pearson's product-moment correlation was used for statistical analyses.

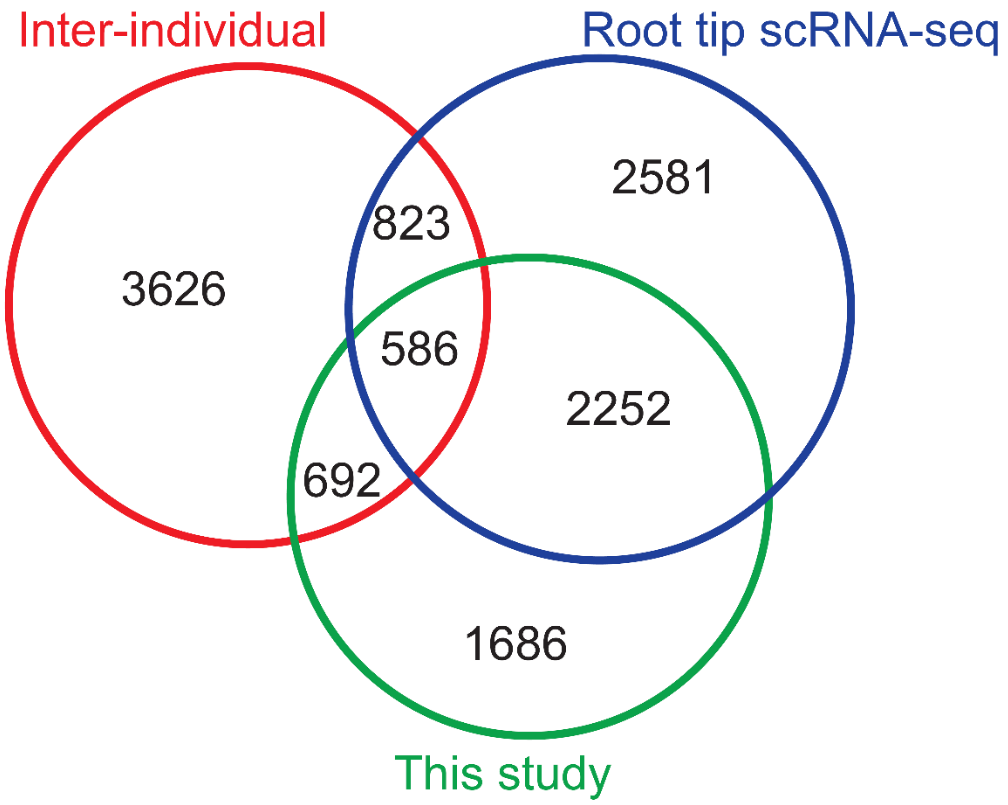


**Fig. S2 | Overlap of lowly variable genes identified from different datasets.**

Venn diagram of lowly variable genes (LVGs) found from different datasets. Inter-individual LVGS are as identified in the study of Cortijo et al (doi: 10.15252/msb.20188591). Root tip scRNA-seq data was obtained from the work of Zhang et al (doi: 10.1016/j.molp.2019.04.004).

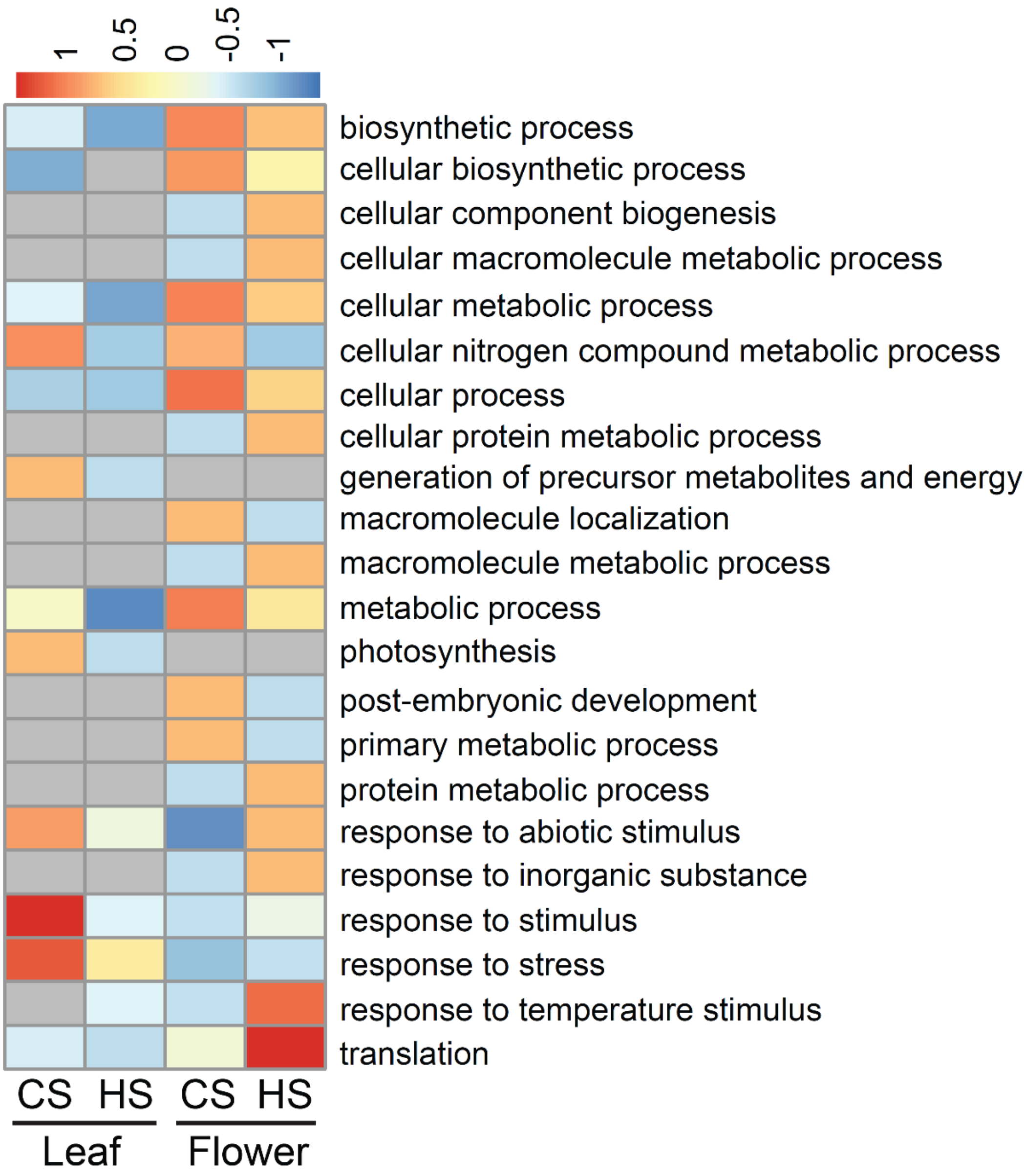


**Fig. S3 | Enrichment of genes associated with low expression variability.**

GO analyses of LVGs identified from each sample indicated.

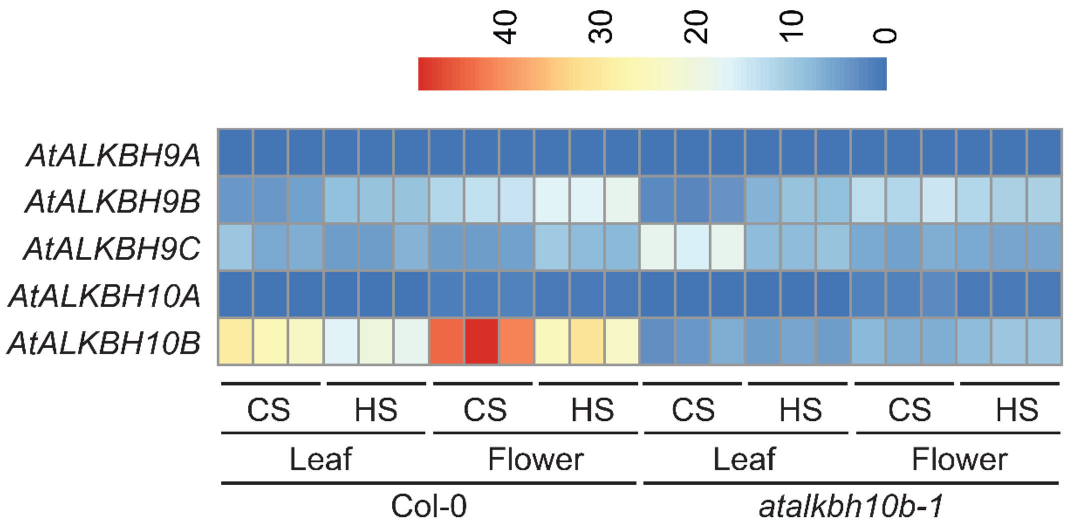


**Fig. S4 | Expression profiling of genes encoding for RNA demethylases.**

Gene expression profile of five RNA demethylases in *Arabidopsis*.

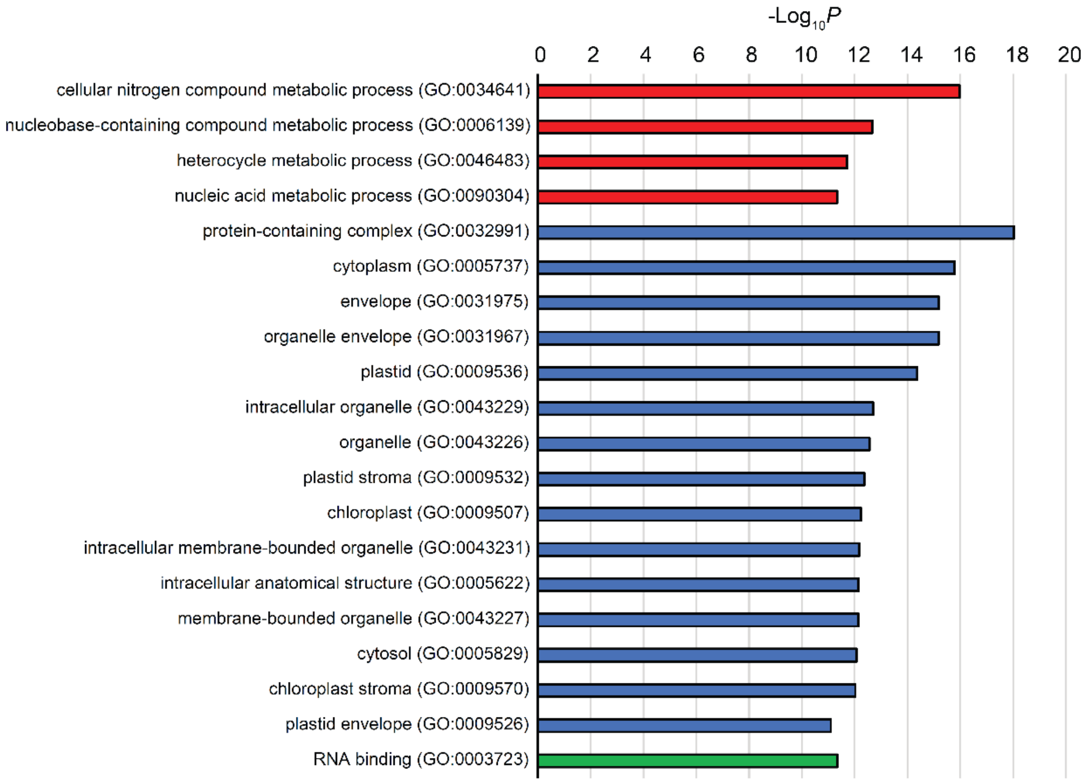


**Fig. S5 | Gene enrichment of hypo-activated genes in the *10b* mutant.**

GO analyses of *10B* targeted genes. Categories with *P* values below $10^{-10}$ are shown.

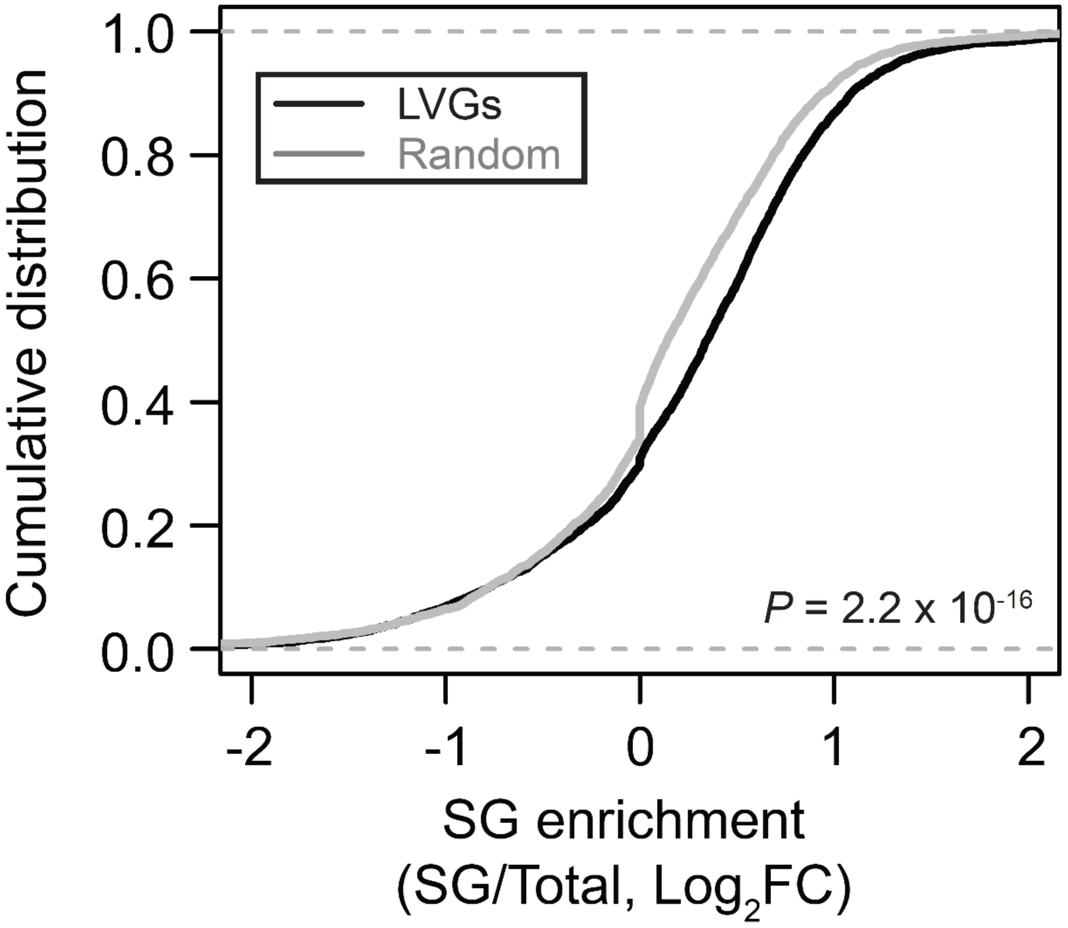


**Fig. S6 | Stress-granule association of lowly variable transcripts.**

SG enrichment score of lowly variable genes in comparison with randomly selected genes. SG enrichment data was obtained from the study of Kim et al (doi: 10.1038/s41477-021-00867-4).
